# Systematic evaluation of isoform function in literature reports of alternative splicing

**DOI:** 10.1101/303412

**Authors:** Shamsuddin A. Bhuiyan, Sophia Ly, Minh Phan, Brandon Huntington, Ellie Hogan, Chao Chun Liu, James Liu, Paul Pavlidis

## Abstract

Although most mammalian genes have multiple isoforms, an ongoing debate is whether these isoforms are all functional as well as the extent to which they increase the genome’s functional repertoire. To ground this debate in data, we established a curation framework for evaluating experimental evidence of functionally distinct splice isoforms (FDSIs) and analyzed splice isoform function for over 700 human and mouse genes. Despite our bias towards prominently studied genes, we found experimental evidence meeting the classical definition for functionally distinct isoforms for only ~5% of the curated genes. If we relax our criteria, the fraction of genes with support for FDSIs remains low (~13%). We provide evidence that this picture will not change substantially with further curation. Furthermore, many FDSIs did not trace to a specific isoform in Ensembl. Our work has implications for computational analyses of alternative splicing and should help shape research around the role of splicing on gene function from presuming large general effects to acknowledging the need for stronger experimental evidence.

## INTRODUCTION

An ongoing debate is whether most mammalian genes produce more than one functional isoform (Blencowe, 2017; Tress *et al*, 2017a, 2017b). The mere presence of multiple isoforms in public sequence databases is clearly insufficient to settle the question (Light & Elofsson, 2013). Arguments against widespread functional alternative isoforms include the fact that the splicing machinery’s limited fidelity causes the stochastic generation of “junk” isoforms (Hsu & Hertel, 2009; Melamud & Moult, 2009). Analyses using proteomics and molecular evolution approaches have also failed to support the expression and conservation of most splice isoforms (Abascal *et al*, 2015b; Pickrell *et al*, 2010; Reyes *et al*, 2013; Saudemont *et al*, 2017; Tress *et al*, 2017b). Nevertheless, the question lingers because the lack of evidence is not generally accepted as evidence and the function of most splice isoforms remain unknown (Blencowe, 2017; Light & Elofsson, 2013). Beyond the question of whether most genes have more than one functional isoform is a critical issue: whether these isoforms increase the functional repertoire of genes, or are merely functionally redundant (Lipscombe *et al*, 2013; Kriventseva *et al*, 2003; Stetefeld & Ruegg, 2005; Lipscombe *et al*, 2002). In this paper we take steps to address the gap between the commonplace assumption that most genes have more than one distinct functional product and the evidence-based reality.

Establishing whether a gene has functionally distinct isoforms requires experimental validation. While databases that contain information on transcript isoforms gather information on isoform features, none attempt to assess functionally distinct isoform reports from the experimental literature. For example, Ensembl, RefSeq, and UniProt catalog and annotate splice isoforms based on evidence that they exist as a transcript or protein (Aken *et al*, 2016; Bely *et al*, 2010; Zhao & Zhang, 2015; Light & Elofsson, 2013). However, the existence of a splice isoform alone does not provide direct support for its functionality, much less functional distinctness.

To establish the extent to which splice isoforms increase the functional repertoire of the genome, we need data on whether genes have *functionally distinct splice isoforms (FDSIs)*. Identification of genes with FDSIs requires experimental support to demonstrate the necessity of each splice isoform. A classical method to determine the function of a given gene is to knock it out and observe the phenotypic consequence (Alberts *et al*, 2002; Shehu *et al*, 2016). This idea readily extends to isoforms; if a single isoform is made absent and that isoform is necessary for the normal function of the gene, then a consequence (change in phenotype) would be expected. A gene has FDSIs if two or more isoforms meet this criterion independently (Figure 1A). In contrast, the depletion of an unnecessary or redundant splice isoform will not cause a phenotype. Another approach that is often used to probe the function of isoforms is overexpression. However, overexpression is well known to be fraught with interpretational challenges including artifacts so the gold standard is to generate loss-of-function alleles (Gibson *et al*, 2013). Note that a negative result from such experiments is not evidence of a lack of functional distinctness, as it is possible the functional distinction between the isoforms may be eventually discovered. Curating the genes with FDSIs is of obvious importance to evaluate the state of the literature support for the commonplace claim that alternative splicing increases the functional repertoire of the genome.

**Figure 1:**
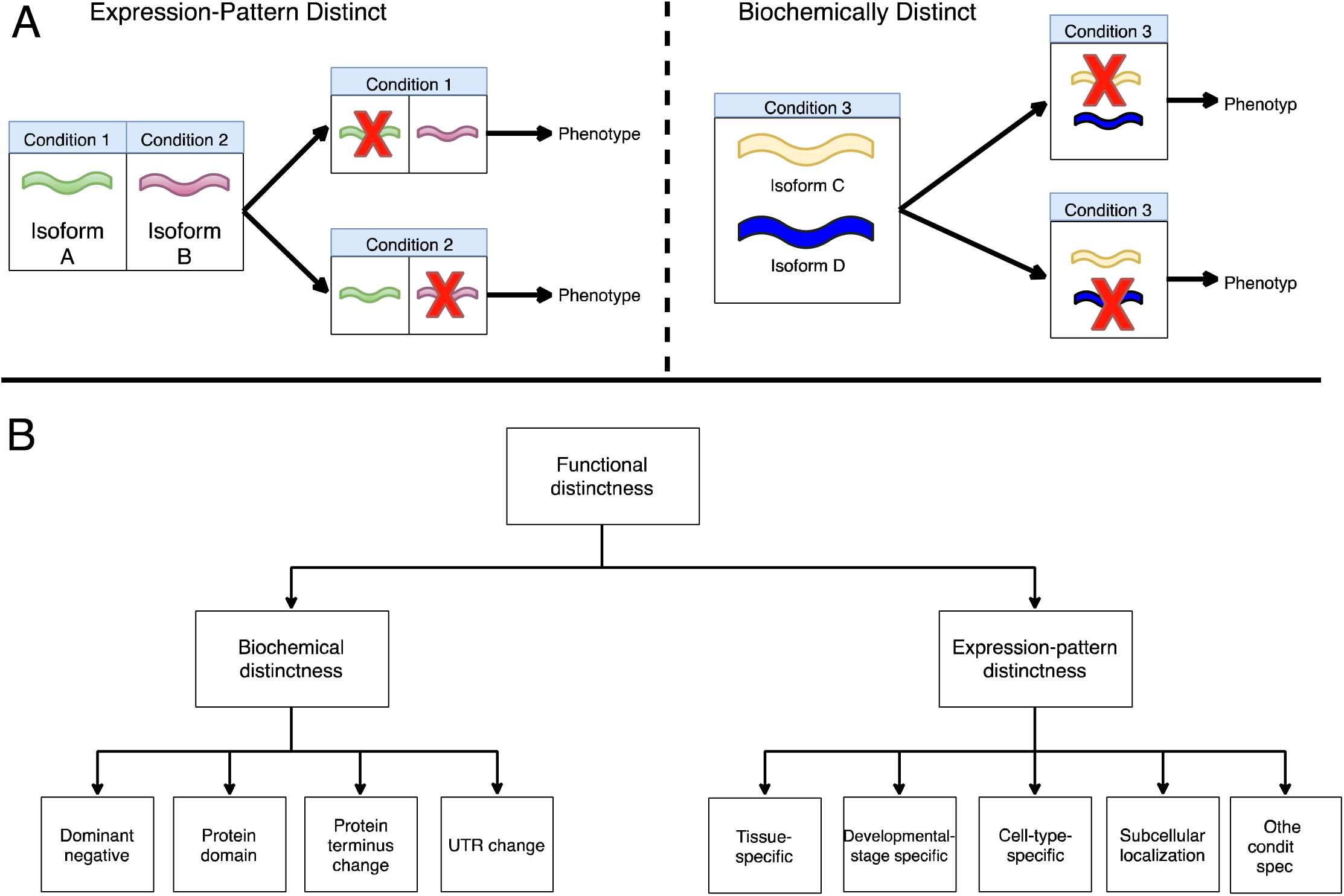
Non-mutually exclusive types of functional distinctness for literature reported genes with FDSIs. **A)** Generally, the distinctness of FDSIs of the same gene can be attributed to expression-pattern distinctness or biochemical distinctness. Expression-pattern distinctness is defined as a gene having specific splice isoforms necessary in distinct conditions. The depletion of the splice isoform in its distinct condition causes a phenotype. Biochemical distinctness is defined as a protein structure difference between splice isoforms of the same genes. While the FDSIs of the gene can be expressed in the same condition, the depletion of either splice isoform causes a phenotype. **B)** For genes with FDSIs, we categorized the specific subtypes of functional distinctness which contributed to the distinctness between the splice isoforms of the gene (summarized in Table 4). Expression-pattern distinctness can be further categorized as “cell-type-specific”, “tissue-specific”, “developmental-stage-specific”, “subcellular localization-specific” and “other condition-specific”. Biochemical distinctness can be further categorized as “dominant-negative”, “protein domain”, “UTR change” and “protein terminus change”.

Beyond identifying knowledge gaps, establishing a set of genes with FDSIs provides potential avenues for improving computational approaches to analyzing alternative splicing. For example, classifiers, such as PULSE, attempt to predict genes with multiple functional splice isoforms (Hao *et al*, 2015). Hao et al. trained PULSE using a set of splice isoforms confirmed by Western blot experiments. PULSE predicted that one-third of human protein-coding genes have multiple functional isoforms (not necessarily functionally distinct). A difficulty cited by Hao et al. was in the identification of training data, an issue which is even worse if one is interested in functional distinctness. Having lists of experimentally validated genes with FDSIs could open the door to improved algorithmic approaches in characterizing isoform function.

Here we present a literature-based analysis of experimental evidence for functionally distinct splice isoforms (FDSIs) for over 700 human and mouse genes. Despite a gene selection strategy that was highly biased towards genes suggested to have multiple functional isoforms, we found good experimental evidence for FDSIs for fewer than 10% of genes.

## RESULTS

### Landscape of the alternative splicing literature

To generate a starting set of papers to curate, we queried PubMed on August 2017 using the term “alternative splicing”. We found 19,049 human studies and 8,197 mouse studies representing 12,891 human genes and 7,585 mouse genes. While the median number of papers per gene was one, there was a large variance (Figure S1 and Figure S2). Most human genes (7,738) had only one such paper associated with them, while some have up to 100 (for example, SRSF1). We also observed that genes with many “alternative splicing”-mentioning papers tend to have many papers in PubMed overall (Spearman’s rank correlation = 0.55, Figure S3). For example, we identified 86 studies linked to human *TP53* with the term “alternative splicing” (rank 2), but this is not particularly remarkable because overall, PubMed contains 8,261 studies linked to TP53 – the most studies for any single gene. This suggests, unsurprisingly, that heavily studied genes tend to have more research done on their splicing.

### Curation Summary

We manually curated primary studies which provide evidence for the function of splice isoforms. As described in Methods, we selected genes and publications for curation in a manner that we expected should enrich for documentation of functional distinctness – for example, using review articles on splicing function. The curation process primarily focused on determining whether the elimination of expression of each splice isoform from a single gene caused an observable phenotype. Table 1 provides a summary of the knowledgebase as of January 20^th^, 2018, and File S1 and S2 contains a list of all curated studies. In total, we curated 1,040 human and mouse studies. This encompasses 829 human studies (528 genes) and 246 mouse studies (206 genes). On average, we have curated a median of 1 study per curated human gene and 1 study per curated mouse gene (mean = 1.5 studies and 1.2 studies, respectively). Our curation evaluations (see Methods) revealed that the curators agree on the interpretation of a paper 98% of the time. Errors were generally false positives for functional distinctness, which we addressed in the final review (see Methods).

**Table 1:**
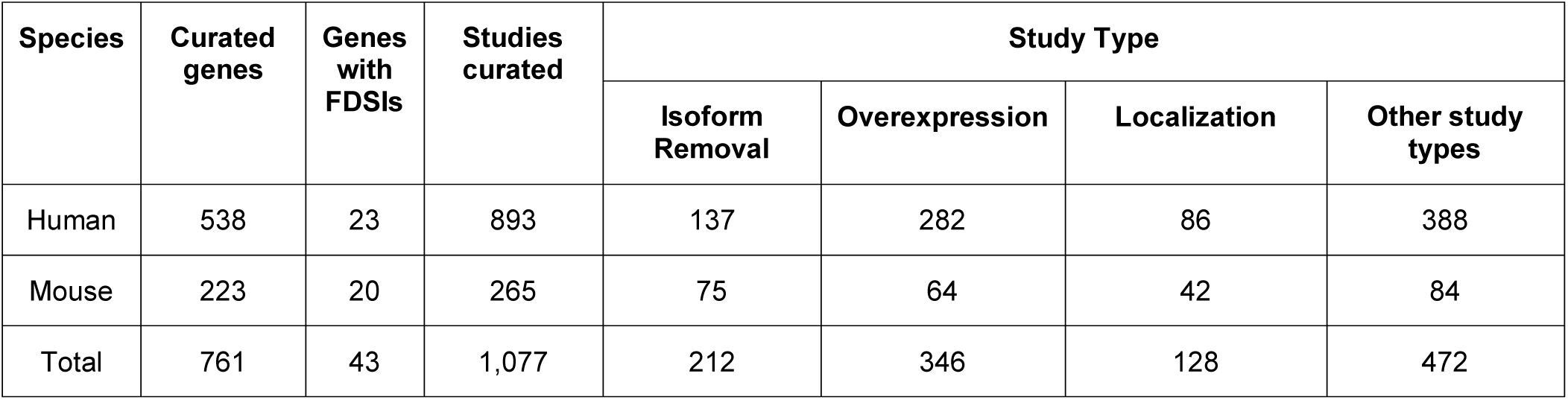
Curation of alternative splicing literature has reveals 23 human genes and 20 mouse genes with functionally distinct splice isoforms (FDSIs). The 23 human genes with FDSIs accounted for almost 4% of human genes annotated in this knowledgebase, while the 20 mouse genes accounted for 9% of the all mouse genes annotated. The majority of curated studies could be classified into three different types: “isoform removal”, “overexpression” and “localization”. Isoform removal studies have experiments where expression of at least one splice isoform is eliminated and a phenotypic change is evaluated. Overexpression studies have experiments where at least one splice isoform is overexpressed. This “abundance” of the splice isoform can cause a phenotype (not necessarily distinct). Localization studies have experiments that characterize where in the cell or organism the splice isoform is expressed. A single study can report experiments with multiple study types. The total number of human and mouse studies curated do not sum to 1,158 studies because some publications investigated both human and mouse forms of a single gene.

| Species | Curated genes | Genes with FDSIs | Studies curated | Study Type |  |  |  |
| --- | --- | --- | --- | --- | --- | --- | --- |
|  |  |  |  | Isoform Removal | Overexpression | Localization | Other study types |
| Human | 538 | 23 | 893 | 137 | 282 | 86 | 388 |
| Mouse | 223 | 20 | 265 | 75 | 64 | 42 | 84 |
| Total | 761 | 43 | 1,077 | 212 | 346 | 128 | 472 |

### Identification of 23 human genes with direct evidence of functionally distinct splice isoforms

By definition, a gene with functionally distinct splice isoforms (FDSIs) has at least two splice isoforms necessary for the gene’s normal function. We find that genes with such evidence are rare: about 4% of curated human genes (9% of mouse genes) have FDSIs, based on reports in a total of 69 studies out of 1,040 studies. Note that 124 studies depleted only one splice isoform of a gene and no other study we curated had depleted any other isoforms of the same gene. We provided the full list of the 23 human genes and the 20 mouse genes with FDSIs in Table 2 with additional information in File S3. RNAi knockdown experiments provided support for over 75% of these FDSIs, while the remaining FDSIs were characterized using gene knockouts combined with isoform-specific rescue.

**Table 2:**
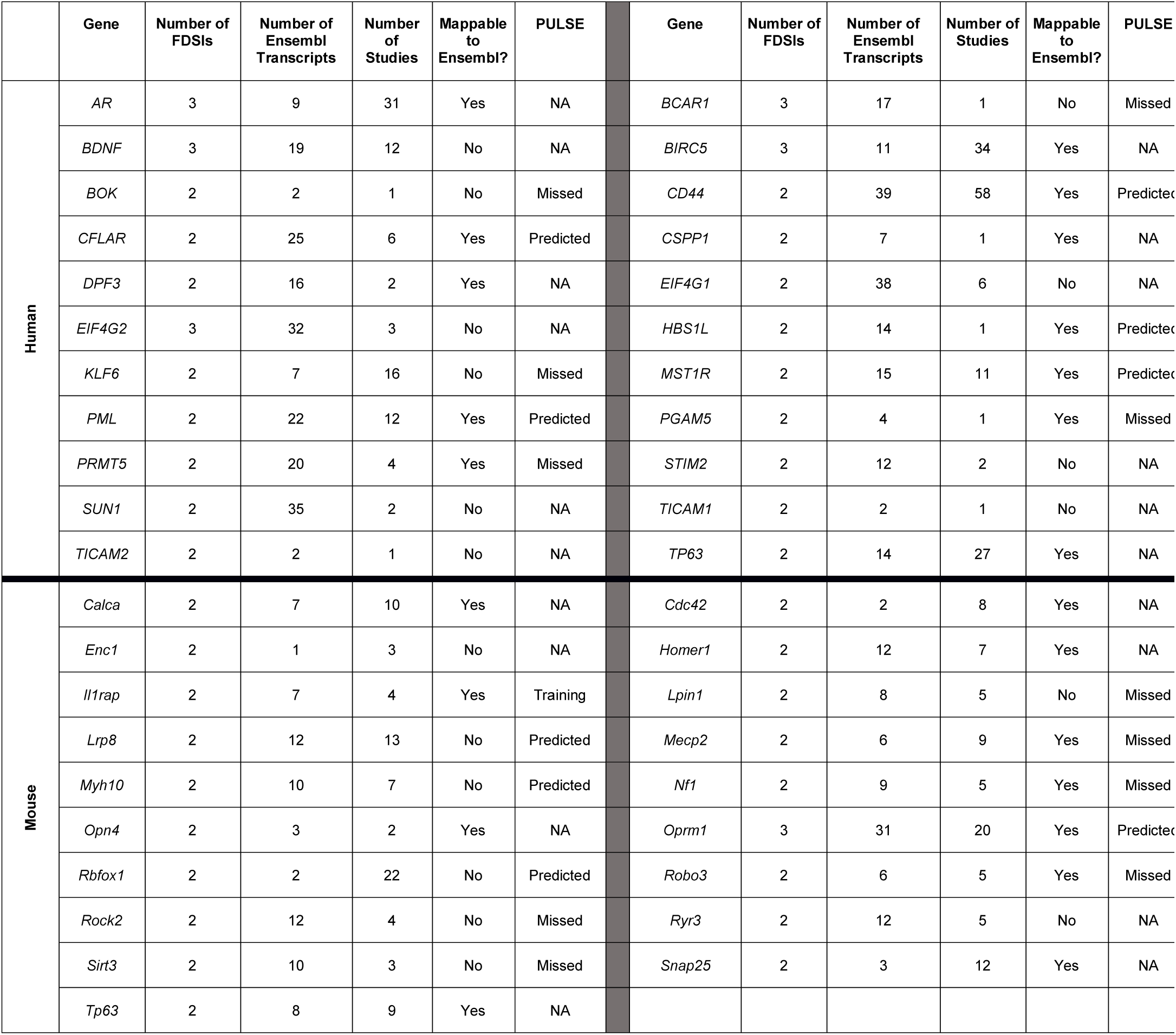
Genes with FDSIs identified. Studies have provided positive evidence of functional distinctness for these genes in experiments where individual splice isoforms were eliminated and a phenotypic change was observed. See File S3 for study demonstrating functional distinctness. “Number of FDSIs” indicates the number of splice isoforms where depletion of splice isoforms causes a phenotype. “Number of Ensembl Transcripts” indicates number of transcripts found in Ensembl entry for gene. “Number of studies” indicates the number of studies associated with the gene retrieved with the term “alternative splicing” on PubMed. The highest number of FDSIs found in a single gene is three. “Mappable to Ensembl” indicates genes where we successfully linked all FDSIs back to Ensembl. “PULSE” indicates whether the gene was used at all by Hao and colleagues in their computational predictions. “Training” in this column means that the gene was used as part of PULSE’s training set. “Predicted” means that PULSE predicted that the gene has multiple functional splice isoforms. “Missed” means that PULSE failed to predict that the gene has multiple functional splice isoforms. “NA” means that the gene was not an input for PULSE.

We sought genes with negative evidence for FDSIs. For these cases, experiments individually depleted multiple splice isoforms for a single gene, however, only one splice isoform’s depletion caused a phenotype and while the depletion of the other splice isoforms caused no phenotype. We found 17 genes with such evidence (shown in Table 3).

**Table 3:** Genes with evidence failing to support FDSIs (negative results). These genes had multiple isoforms tested however only one splice isoform caused a change in phenotype.

| Gene | Experimental method | Tissue/Cell Type | Reference (PubMed ID) |
| --- | --- | --- | --- |
| *Ank3* | Isoform-specific rescue | Neuron | 25552556 |
| *Ar* | Knockdown | Prostate cancer cell line | 20823238 |
| *Ccnd1* | Isoform-specific rescue | Embryonic fibroblast | 21200149 |
| *Dab1* | Isoform-specific rescue | Neuron | 28968791 |
| *Dntt* | Isoform-specific rescue | Bone marrow | 11136823 |
| *FANCE* | Isoform-specific rescue | Breast cancer cell line | 26277624 |
| *FNBP1L* | Isoform-specific rescue | MDCK cell line | 26063734 |
| *MADD* | Isoform-specific rescue | Kidney | 16682944 |
| *Pcdha1* | Isoform-specific rescue | Brain | 18973563 |
| *PDE4D* | Knockdown | Kidney | 16030021,<br>17673687 |
| *PEX19* | Isoform-specific rescue | Fibroblast | 11883941 |
| *Pparg* | Knockdown and isoform-specific rescue | Adipose | 11782442 |
| *RAP1GSDS1* | Knockdown | Breast cancer cell line | 24197117 |
| *RREB1* | Knockdown | Bladder | 21703425 |
| *SIRT1* | Knockdown | Colon cancer cell line | 22124156 |
| *Smad2* | Isoform-specific rescue | Embryonic stem cells | 15630024 |
| *STAT1* | Knockdown | Embryonic cells | 21914475 |

As mentioned, we biased our gene and paper selection in such a way that our estimate of 4% (9% for mouse) might be too high. To help clarify this issue, we randomly selected 100 human genes (from those that had at least one alternative splicing related paper) for gene-centric curation (listed in Table S1). Of these 100 genes, two genes (PML and DPF3, 2%, of the curated genes) had experimental evidence of FDSIs.

We also curated gain-of-function experiments where investigators overexpressed multiple splice isoforms of the same gene. From our 538 curated human genes and 223 curated mouse genes, we found 47 human genes (~9%) and 10 mouse genes (~4%) where investigators overexpressed individual splice isoforms and yielded multiple distinct phenotypes. Such studies did not meet our criteria for FDSIs, but we report them in case this relaxed criterion is of interest to others.

### Genes tend to express functionally distinct splice isoforms in the same condition

To further explore functional distinctness in splicing, we identified non-mutually exclusive types of functional distinctness between FDSIs of the same gene, summarized in Table 4. We classified two main types of distinctness, expression-pattern distinctness and biochemical distinctness. Genes with expression-pattern distinct FDSIs have splice isoforms necessary for specific conditions while genes with biochemical distinctness have FDSIs with distinct biochemical properties that cannot compensate for each other even when co-expressed (for further description see Methods and Figure 1).

**Table 4:**
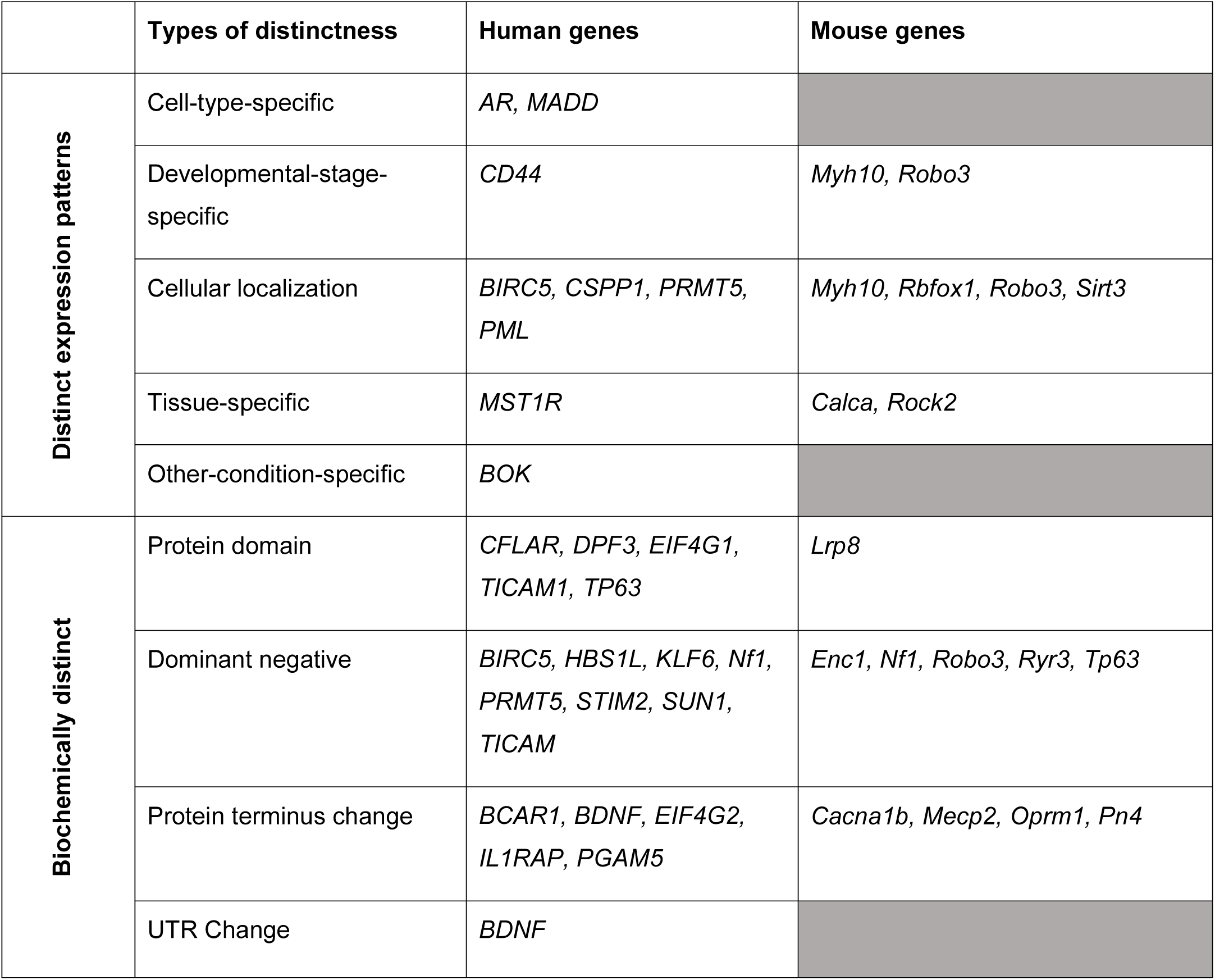
Most genes with FDSIs have biochemically distinct splice isoforms. Genes with FDSIs were categorized on functional type based on the literature that reported on the FDSIs using the scheme outlined in Figure 1. Genes categorized as “distinct expression patterns” express FDSIs in specific conditions. Genes categorized as “biochemically distinct” have FDSIs whose functional distinctness is a consequence of biochemical differences in their final protein product. Genes can be categorized as both “distinct expression patterns” and “biochemically distinct” such as Myh10 and Robo3.

The majority of genes (27/43) have biochemically distinct isoforms, rather than expression-pattern distinct. We identified “dominant-negative” as the most common subtype of biochemically distinct FDSIs (12/31 genes). For example, the mouse gene *Enc1* has two FDSIs, named “57 kDa” and “67 kDa” by the authors, interacting in the Wnt-signalling pathway (Worton *et al*, 2017). Knockdown of 57 kDa promoted osteoblast mineralization while the knockdown of 67 kDa inhibited osteoblast mineralization.

In contrast to biochemical distinctness, we identified fewer cases of genes with expression-pattern distinct FDSIs. Only a total of 17 human and mouse genes had FDSIs in which the distinctness arises from distinct expression patterns. For example, the mouse gene *Myh10* has two FDSIs, named B1 and B2 by the authors (Ma *et al*, 2006). Cells in the brainstem express B1 to promote normal migration of facial neurons, while cells in the cerebellum expressed B2 to promote normal cerebellar Purkinje cell development.

### Challenges linking FDSIs to sequence databases

We attempted to link all identified FDSIs back to Ensembl transcript identifiers and were successful in 25/43 cases. Our process was as follows. First, in the studies for seven genes, investigators provided a GenBank or RefSeq ID. We were able to map three of these to Ensembl (which includes GenBank and RefSeq data), but not for the other four (for more details see File S3), accounting for four of the 16 failures. Next, for 36 genes with missing accession information, we used sequence alignment or other information to identify likely matches (See Methods). This was successful in 25 cases. In a further 6 cases, we were able to determine a sequence by referring to other papers by the same authors. Despite extensive efforts, we were unable to find matching Ensembl transcripts for the isoforms of 5 genes. This situation was not specific to Ensembl as we failed to link isoforms of 8 genes to UniProt; see File S3.

### Only a quarter of genes with FDSIs are predicted by a computational classifier

Hao et al. (Hao *et al*, 2015) developed a machine learning algorithm (PULSE) that predicted 1/3 of human genes have more than one functional isoform. We hypothesized that our curated genes with FDSIs would be enriched among those predictions, because even though Hao and colleagues were not attempting to predict functional distinctness, genes with FDSIs by definition have more than one functional isoform. Though we included PULSE’s training genes in our gene-centric curation, only two gene with FDSIs (including human orthologues of our curated mouse genes) were used by Hao et al., in their training data. In their validation gene set of 212 genes, we found none of our genes with FDSIs. Hao et al. predicted 2,419 genes to each have multiple functional splice isoforms. Ten of our genes with FDSIs are included in this set. Based on input set used for PULSE predictions, the classifier failed to predict 12 of our genes with experimentally-validated FDSIs to have multiple functional splice isoforms. However, our interpretation of these is limited because of the small number of genes with FDSIs.

## DISCUSSION

This paper represents progress towards documenting and evaluating the breadth of evidence for functionally distinct splice isoforms (FDSIs) for human and mouse genes. The inspiration for our study was strong arguments against the likelihood of most genes having multiple functional isoforms, contrasted with the ubiquitous claim that splicing vastly increases the functional repertoire of the genome (Tress *et al*, 2017b; Wang *et al*, 2008a; Lipscombe *et al*, 2013; Kriventseva *et al*, 2003; Lipscombe *et al*, 2002; Stetefeld & Ruegg, 2005; Auboeuf, 2018). This led us to ask where this latter claim comes from: while surely there are interesting cases of multi-isoform genes, has this been optimistically extrapolated to the entire genome? Our analysis suggests this is the case and supports the hypothesis that the majority of splice isoform functions remain unknown (Light & Elofsson, 2013; Frankish *et al*, 2012; Mudge *et al*, 2011). While we were not surprised that there is no evidence of FDSIs for most genes, we were surprised by the low fraction for which there is supporting data, a mere 4% in human genes and 9% in mouse genes. Regardless of whether this number holds true with more curated studies, by contributing a list of genes with documented functionally distinct isoforms, we start to identify the scope of the gaps, the parameters for future experimental work, and assist computational methods that require training examples.

The low fraction of genes surveyed for which we found evidence of FDSIs (~4–9%) agrees with the general sense that we still have limited concrete evidence of more than one functional splice isoform per gene (Kelemen *et al*, 2013; Reyes *et al*, 2013; Tress *et al*, 2017b). Even if we loosen our criteria to include overexpression studies, this fraction rises only to ~12–13% Furthermore, we only considered genes for which some literature exists for their isoforms, so the range 4% to 9% is relative to genes that have at least one publication about them associated with splicing. Based on our PubMed queries, we estimate that one-third of human protein-coding genes do not have any type of specific experimental study of differences among their isoforms. For most genes the main available sources of information come from genome-wide studies of transcript expression patterns, which do not address function.

One might question whether the fraction 4% will rise substantially as we continue our curation efforts, but we hypothesize a lower true fraction of genes with documentation of FDSIs in the literature. First, we aimed the gene-centric aspect of our curation at genes mentioned in review articles or otherwise prominent genes, and thus is highly biased towards genes with experimentally-backed function, yielding an over-estimate. Second, the gene-centric survey of 100 randomly-selected human genes yielded only two genes with evidence of FDSIs. Third, we found a median of only one study per gene from PubMed. Since the genes with FDSIs tended to be genes with relatively more associated studies (Table 2), genes with few associated studies seem less likely to yield existing positive evidence for functional distinctness. Fourth, investigators face technical and/or resource challenges when testing the functional distinctness of isoforms, requiring either the ability to conduct isoform-specific depletion experiments, or isoform-specific rescue following a complete gene knockout. Reasonably, one might suppose that in many cases the experiments have not been done. The essential problem remains that most genes simply have not had their isoforms tested in such a way as to establish distinct functions.

We also sought *negative* evidence of genes having FDSIs from experiments where the depletion of only one splice isoform causes a phenotype while the depletion of the remaining splice isoforms of the same gene causes no phenotype. However, we only identified eight human genes and nine mouse genes from 17 studies with this type of evidence in our current curation of 1,077 studies (Table 3). Since most studies consider only one type of functional assay, it remains possible that tests of different functions would yield positive results for these genes. Nevertheless, the “file-drawer effect” – a type of publication bias against negative results – potentially plays a role in the dearth of negative evidence (Kennedy, 2004).

### Evaluating the evidence for FDSIs at the gene level

After the curation of over 1,000 alternative splicing studies, we identified 22 human genes and 20 mouse genes with evidence for functionally distinct splice isoforms, mostly determined by RNAi knockdown experiments. RNAi knockdowns naturally align with our definition of a functional splice isoform and how researchers traditionally determine function in molecular biology. One question that arises in discussing RNAi is target specificity and efficacy. In most, but not all, of the papers we curated as having FDSIs, the authors demonstrate the target specificity of their siRNA to effectively deplete a single isoform. We raise this as a reminder that reports of evidence for functional distinctness may vary in quality.

Isoform-specific rescues demonstrating functional distinctness provide an alternative option to knockdown studies but the method has limitations when determining whether the splice isoforms rescue distinct phenotypes. In some studies, splice isoforms of the same gene clearly rescued distinct functions. For example, Candi and colleagues performed rescue experiments on Tp63-null mice (Candi *et al*, 2006). The knockout of Tp63 impeded the development of skin. In the rescue experiments, the splice isoform ΔNp63 restored the skin’s basal layer while the TAp63 restored the skin’s upper layers. In contrast, other studies rescued the same phenotype with each splice isoform, which makes evidence of functional distinctness unclear. For example, in the investigation by Coldwell and colleagues, each splice isoform of *EIF4G1* (eIF4G1e and eIF4G1f) rescued the phenotype of translation by restoring the translation rate (Coldwell *et al*, 2012). It is unclear whether this constitutes evidence of functional distinctness. Since both splice isoforms rescued the same phenotype, they appear functionally redundant. Nevertheless, in cases such as these, we accepted the claim of the authors that the gene has FDSIs.

We resisted accepting overexpression studies as demonstrating FDSIs for two reasons. First, overexpression experiments are known to be subject to a variety of artifacts (Gibson *et al*, 2013). Second, and more importantly, overexpression experiments fail to provide evidence for a splice isoform’s necessity. In molecular biology, a molecule’s necessity can only be supported by the effects of the molecule’s absence (Gannett, 1999; Gifford, 1990). Thus, we have more confidence in isoform depletion experiments to provide support for genes with FDSIs compared to overexpression. We draw a parallel to the standards of evidence for characterizing gene function, in which evaluation of a loss of function is the gold standard (Kopp & Mendell, 2018). We argue that the same criteria used to establish gene function must be applied to isoforms.

### Types of functional distinctness in FDSIs

It has been speculated that many poorly-characterized isoforms may have function because genes express splice isoforms in specific conditions, perhaps yet to be studied (Blencowe, 2017; Pan *et al*, 2008; Wang *et al*, 2008b). It is therefore relevant that the minority (17) of genes had functional distinctness due to condition-specificity. This may simply be due to a lack of study of condition-specific studies, as it might be generally easier to study isoforms expressed in the same conditions. Our results thus point to a potential gap in the literature.

### Disconnect between the literature and gene databases

In one-third of the genes with FDSIs, the isoforms studied in a paper could not be matched to transcripts in Ensembl (as mentioned, this is not an Ensembl-specific problem; ~20% of genes had functional isoforms that could not be matched to UniProt). Conversely, Ensembl contains many transcripts that the literature ignores. This observation has fairly serious implications for basing splice isoform research on the contents of Ensembl (or related databases). If one developed experiments to functionally test the splice isoforms of the genes we identified to have FDSIs based on Ensembl transcripts for that gene, their experiments would not contain the correct FDSIs in at least one-third of the genes. In bioinformatics research, computational methods that make predictions based on Ensembl transcripts might be valueless to experimental biologists as Ensembl does not reflect the literature. Large-scale databases specialized for alternative splicing, such as the Alternative Splicing encyclopedia (ASpedia) and the APPRIS database, tend to anchor to Ensembl (Hyung *et al*, 2018; Rodriguez *et al*, 2018). Of note, previous discussions used APPRIS to understand the functional impact of alternative splicing (Tress *et al*, 2017b). The disconnect between Ensembl the literature also impacts datasets not specific to splicing but where splice isoform information is important. For example, the GTEx consortium provides transcript-level quantification based on the Ensembl transcriptome (Lonsdale *et al*, 2013). The FDSIs that are not in Ensembl are therefore not included in GTEx. Given the few known cases of genes with FDSIs and PULSE’s inability to predict all our genes with FDSIs, it remains crucial that computational resources contain FDSIs and experimentalists ensure that they submit their sequence data to these resources.

### Implications for alternative splicing’s impact on gene function

Recent studies have challenged whether *most* genes can produce multiple functional splice isoforms and our results can offer something to both sides of the debate. We acknowledge that other researchers may have different definitions of a functional splice isoform, but we view the debate within our operational definition – a functional splice isoform is one that is necessary for the gene’s overall function.

On one side of the debate claims that most genes have multiple functionally distinct isoforms and therefore (Blencowe, 2017). Viewing our findings optimistically, we provide what is to our knowledge the only substantial list of human and mouse genes for which this is actually documented to be true. The low number of genes with such evidence can be interpreted as a vast opportunity for experimentalists to identify the functions of the isoforms for >80% of genes. The other side of the debate approaches alternative splicing with a less Panglossian view, with the null hypothesis being that most isoforms do *not* have a specific distinct function (Gould & Lewontin, 1979). Multiple studies taking a genomic or evolutionary perspective have concluded that it is unlikely that most genes have multiple functional splice isoforms (Hu *et al*, 2017; Melamud & Moult, 2009; Hsu & Hertel, 2009; Reyes *et al*, 2013; Abascal *et al*, 2015a; Tress *et al*, 2017b; Light & Elofsson, 2013; Saudemont *et al*, 2017; Kurmangaliyev & Gelfand, 2008; Wang *et al*, 2014; Zhang *et al*, 2009; Pickrell *et al*, 2010). Viewed pessimistically, our data is consistent with this body of work. If the literature lacks supporting evidence for widespread FDSIs, the null hypothesis should be maintained and claims that every observed isoform has a function to be discovered should be viewed skeptically.

### Conclusions

To our knowledge, this report represents the first effort to curate the literature in order to determine the genes where splicing increases the genome’s functional potential. Such individual reports have been generally ignored in the debate about the function of alternative splicing, which has instead focused on databases and high-throughput data sets. Our estimate that only 4% of human and 9% of mouse genes have evidence for functionally distinct isoforms serves both a sobering reminder of the limited evidence, and a motivation for increased experimental efforts to settle the debate. At the same time, we also recognize there are likely genes with FDSIs that we did not curate and should be included. We invite contributions from the community of corrections or suggestions for papers or genes to curate, which can be sent to the authors.

## MATERIAL AND METHODS

### Determining the type of functional distinctness

We developed a scheme to describe non-mutually exclusive types of functional distinctness found in genes with FDSIs. We recognize two general biological mechanisms by which functional distinctness could arise, schematized in Figure 1A, and elaborated on further below: “expression-pattern distinctness” or “biochemical distinctness”. Figure 1B outlines our full scheme for classifying functional distinctness. The subclasses we identified were designed to accommodate how functional distinctness is reported in the literature we curated, that is, we did not create this classification wholly *ab initio*. We determined the type of functional distinctness using the publication which provided the evidence for FDSI, but some cases required an inference based on other literature by the authors. We stress that a gene can have multiple types of functional distinctness. For example, biochemically distinct isoforms could also have expression pattern distinctness. We annotated as many types of functional distinctness as were provided by the literature reports.

### Expression-pattern distinctness

Expression-pattern distinctness requires the condition-dependent expression of isoforms of a single gene. Generally, in this category, splice isoforms of the same gene have functional relevance in distinct conditions. We further specified expression-pattern distinctness as “subcellular-localization-specific”, “cell-type-specific”, “tissue-specific”, “developmental stage-specific”, and “other-condition-specific”. Thus genes with cell-type-specific FDSIs express their splice isoforms in distinct cell types, and the elimination of expression of either splice isoform causes a phenotype (Figure 1A). Crucially, the splice isoforms’ final products could be identical (that is, they are not biochemically distinct). However, they are still functionally distinct because they have at least partly non-overlapping expression patterns so one cannot fully compensate for the other.

### Biochemical distinctness

Biochemical distinctness is defined as differences in biochemical properties or activities, and which cannot compensate for each other even if co-expressed in the same condition. We further specified biochemical distinctness as “protein domain change”, “dominant negative”, “subcellular localization”, “UTR change” and “protein terminus change”. Genes categorized as FDSIs with distinct protein-domains indicate that each splice isoform has a unique structural or functional unit in their final protein product. We manually extracted information about the specific protein domain from the authors providing the evidence of functional distinctness. In some cases, this could involve the presence or absence of one or more protein domains. Genes categorized as “protein terminus change” indicates that the FDSIs’ final protein product differ from each other either in their C-terminus or their N-terminus. These changes to the C- or N-termini usually do not affect the presence or absence of protein domains (or the paper did not make any note of changes to protein domains). Genes with dominant-negative FDSIs have splice isoforms with antagonistic phenotypes. Typically, these splice isoforms regulate each other’s function. The loss of one splice isoform generally affects the function of the other splice isoform. Gene categorized as “UTR change” indicates that the FDSIs of the same gene differ in the UTRs of the mRNA (coding regions may change as well).

### Literature selection

On July 17^th^ 2017, we generated a “starting set” of publications associated with human and mouse genes to curate using PubMed e-utilities and the search term “alternative splicing”. From here curation was both “gene-centric” and “paper-centric.”

### Gene-centric curation

The gene-centric approach attempted to curate all relevant studies associated with a specific gene. PubMed linked each study from our starting set to a specific gene which provided a list of genes with literature. The genes we selected to curate from this list were genes suggested to us by the community, PULSE’s training genes or commonly discussed by the literature (Hao *et al*, 2015). As suggestions form the community might be biased, 100 random genes were also selected for gene-centric curation.

### Paper-centric curation

The paper-centric approach attempted to curate literature likely enriched for evidence of genes with FDSIs. Using this approach, we make no attempt to curate all relevant reports for any specific gene. As a targeted source of literature likely to be enriched for functional evidence, we used review articles on the function of alternative splicing that provided citations for 603 genes (Kelemen *et al*, 2013; Kovacs *et al*, 2010; Ramanouskaya & Grinev, 2017; Stamm *et al*, 2005; Tress *et al*, 2017b; Lipscombe *et al*, 2013). We further extended paper-centric curation with specific search phrases in PubMed. Search terms were: “functionally distinct splice isoforms”, “CRISPR alternative splicing”, “alternative splicing knockdown” and “alternative splicing knockout.” These queries identified an additional 260 papers for our starting set of papers. The genes found in the publications retrieved by these PubMed queries and provided in the aforementioned reviews further informed us of which genes to gene-centrically curate. For example, *BDNF* and *XBP1* were commonly reviewed in the literature and consequently, we gene-centrically curated them.

### Curation process

For each paper, a trained curator first identified general features of that study by manually extracting the following information: the investigated gene, the reported number of the splice isoforms for the gene, the names used by the authors for the splice isoforms, the number of splice isoforms specifically investigated in the paper (“the investigated isoforms”), the experiments performed, the organism where the gene was identified, the organism or cell line used for the experiments, and any claims of functional distinctness.

Next, using a decision tree (Figure 2), we annotated each paper as to whether the data provided positive evidence of functional distinctness for the investigated splice isoforms. We sought evidence where the loss of one isoform (via knockdown, knockout or other means of isoform-specific depletion) produced a phenotype in the test system. We also curated experiments which performed overexpression analyses, which were retained as a separate category from the isoform loss studies (as an example, see study by Scotton and colleagues (Scotton *et al*, 2006). We did not accept studies of aberrant isoforms caused by rare mutations (for example in cancer), as we deemed these as not relevant to the normal function of the gene as we have defined it (as an example, see Cogan et al. (Cogan *et al*, 2012)). If a study provided evidence where investigators depleted multiple splice isoforms of a single gene but at most one splice isoform caused a phenotype, we classified the gene as having *negative* support for FDSIs. Finally, regardless of study type, the curators provided a concise explanation of the functions investigated.

**Figure 2:**
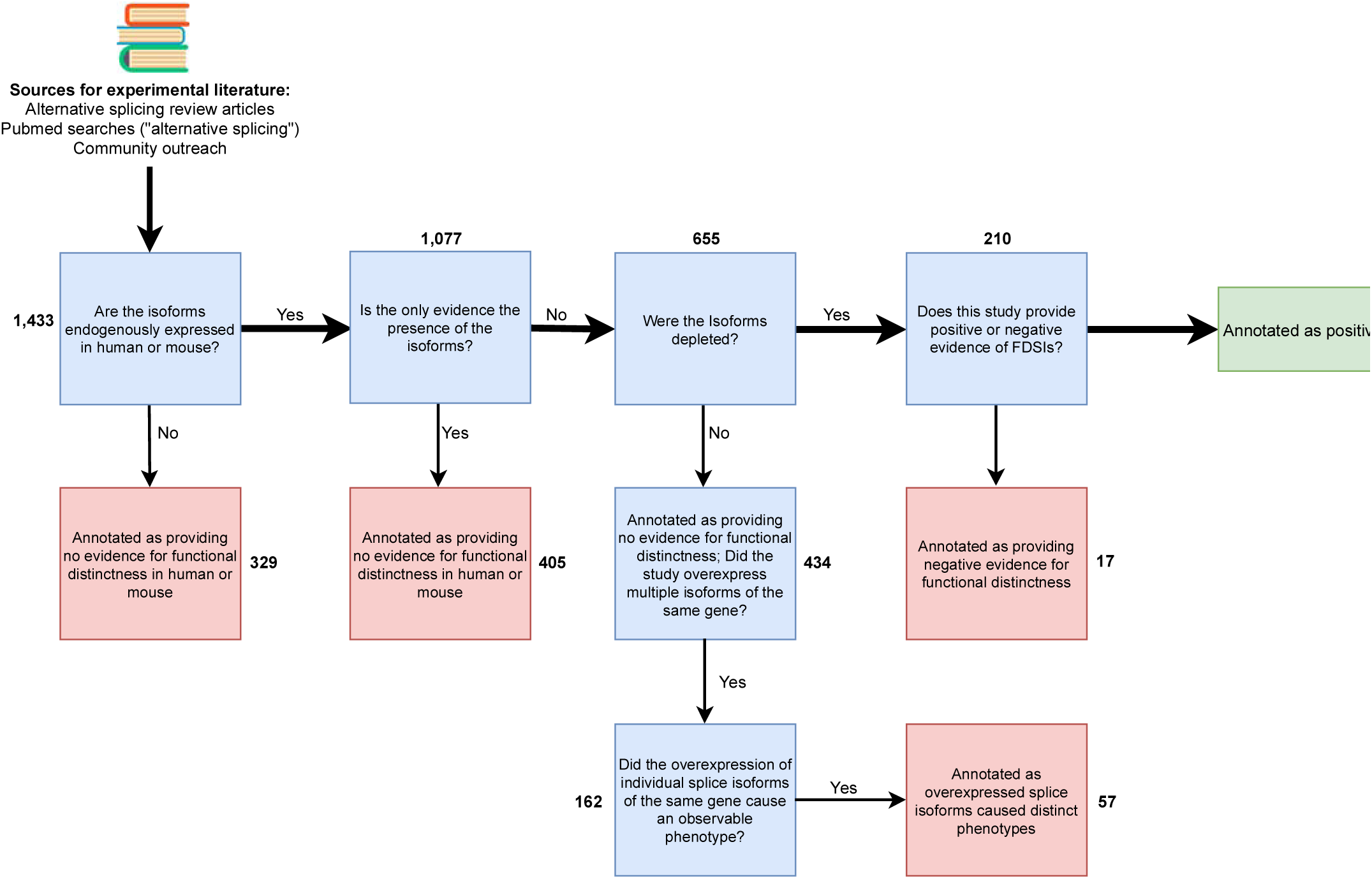
Overview of literature curation scheme. We sought papers which study the functional distinctness of a single human or mouse gene’s splice isoforms. Positive studies are those that provide evidence where multiple splice isoforms of a single gene are depleted and at least two isoforms show a phenotype. We annotated studies as providing negative evidence for functional distinctness when investigators deplete multiple splice isoforms of the same gene but only one produces an observable phenotype. The numbers in bold represent the number of studies in each category.

For our definition of genes with FDSIs, we required evidence for the independent depletion of at least two splice isoforms of the same gene. If the curated study investigated the outcome of the absence of a single isoform for a given gene, then that study alone insufficiently provides evidence of FDSIs. While such studies demonstrate an existence of a single functional isoform, the support for FDSIs requires data on at least two isoforms from the same gene. However, we subsequently attempted to identify a second paper that provided functional evidence for a different splice isoform of the same gene. In situations where a second paper identified evidence of a different functional splice isoform, we recorded the gene as having FDSIs.

### Curator validation

To ensure consistent curation, we evaluated the curators. These tests consisted of all curators curating the exact same randomly selected 50 papers. After the test, we addressed any discrepancy between curators and we updated the curation standards with any necessary clarifications (curation standards provided in File S1). This evaluation process was conducted three times. We also further scrutinized papers annotated as providing positive evidence of a gene with FDSI to eliminate any false positives.

### Linking FDSIs to Ensembl

If a paper provided positive evidence for FDSIs, we linked the splice isoforms with the appropriate Ensembl transcript ID. Generally, studies provided GenBank or RefSeq accession IDs and these accession IDs linked to Ensembl. In the absence of an accession ID, we referred to the literature for sequence information about the splice isoforms and aligned splice isoform sequences to Ensembl using ClustalOmega (Sievers *et al*, 2011).

### Computational predictions of genes with FDSIs

PULSE, a computational classifier developed by Hao et al., predicted 2,419 of 15,639 UniProt genes to have multiple functional isoforms based on a training set of 145 genes (Hao *et al*, 2015). We downloaded the supplementary data provided by Hao et al. to determine whether PULSE predicted our genes with FDSIs to have multiple functional splice isoforms. We also investigated whether any of our genes with FDSIs were part of PULSE training and validation set of genes. This was of interest because a training set enriched for genes with FDSIs may yield predictions for genes with FDSIs, even though PULSE was only designed to detect function, not distinct function. For our comparison to PULSE prediction, we used the human orthologue for any mouse gene with FDSI as determined by BioMart (Smedley *et al*, 2009).

## SUPPLEMENTARY DATA

Supplementary Data are available online.

## AUTHORS’S CONTRIBUTION

SB and PP designed the overall project and curation framework. SB and PP supervised the curators. SL, MP, BH, EL, CL, JL curated the literature. All curators provided feedback to SB and PP for the curation framework. JL queried PubMed for literature to curate. SL referenced splice isoforms to genomic databases. SB determined types of functional distinctness. SB and PP prepared the manuscript.

## ACKNOWLEDGEMENT

The authors would like to thank and acknowledge Dr. Sanja Rogic, Dr. Lilah Toker, Dr. Shreejoy Tripathy and Benjamin Callaghan for their useful feedback throughout the project. We also thank all contributors who provided genes or studies to our curation efforts.

## FUNDING

This work was supported by a Natural Sciences and Engineering Research Council (http://www.nserc-crsng.gc.ca/) CREATE scholarship for the University of British Columbia Bioinformatics Training Program (awarded to SB), and Natural Sciences and Engineering Research Council Discovery grant (RGPIN-2016-05991) and National Institutes of Health (www.nih.gov) grant MH111099 to PP. The funders had no role in study design, data collection and analysis, decision to publish, or preparation of the manuscript.

## CONFLICT OF INTEREST

The authors declare that they have no conflict of interest

## REFERENCES

Abascal F, Ezkurdia I, Rodriguez-Rivas J, Rodriguez JM, Pozo A del, Vázquez J, Valencia A & Tress ML (2015a) Alternatively Spliced Homologous Exons Have Ancient Origins and Are Highly Expressed at the Protein Level. PLOS Comput. Biol. 11: e1004325

Abascal F, Tress ML & Valencia A (2015b) Alternative splicing and co-option of transposable elements: the case of TMPO/LAP2α and ZNF451 in mammals. Bioinformatics 31: 2257–2261

Aken BL, Ayling S, Barrell D, Clarke L, Curwen V, Fairley S, Banet JF, Billis K, Girón CG, Hourlier T, Howe K, Kähäri A, Kokocinski F, Martin FJ, Murphy DN, Nag R, Ruffier M, Schuster M, Tang YA, Vogel J-H, et al (2016) The Ensembl gene annotation system. Database 2016: baw093

Alberts B, Johnson A, Lewis J, Raff M, Roberts K & Walter P (2002) Studying Gene Expression and Function. Available at: https://www.ncbi.nlm.nih.gov/books/NBK26818/ [Accessed April 19, 2017]

Auboeuf D (2018) Alternative mRNA processing sites decrease genetic variability while increasing functional diversity. Transcription 9: 75–87

Bely B, Martin MJ & Apweiler R (2010) Source of annotations in the UniProt Knowledgebase. F1000Posters 1: Available at: http://f1000.com/posters/browse/summary/243 [Accessed May 29, 2014]

Blencowe BJ (2017) The Relationship between Alternative Splicing and Proteomic Complexity. Trends Biochem. Sci. 0: Available at: http://www.cell.com/trends/biochemical-sciences/abstract/S0968-0004(17)30070-1 [Accessed May 9, 2017]

Candi E, Rufini A, Terrinoni A, Dinsdale D, Ranalli M, Paradisi A, De Laurenzi V, Spagnoli LG, Catani MV, Ramadan S, Knight RA & Melino G (2006) Differential roles of p63 isoforms in epidermal development: selective genetic complementation in p63 null mice. Cell Death Differ. 13: 1037–1047

Cogan J, Austin E, Hedges L, Womack B, West J, Loyd J & Hamid R (2012) Role of BMPR2 alternative splicing in heritable pulmonary arterial hypertension penetrance. Circulation 126: 1907–1916

Coldwell MJ, Sack U, Cowan JL, Barrett RM, Vlasak M, Sivakumaran K & Morley SJ (2012) Multiple isoforms of the translation initiation factor eIF4GII are generated via use of alternative promoters, splice sites and a non-canonical initiation codon. Biochem. J. 448: 1–11

Frankish A, Mudge JM, Thomas M & Harrow J (2012) The importance of identifying alternative splicing in vertebrate genome annotation. Database J. Biol. Databases Curation 2012: bas014

Gannett L (1999) What’s in a Cause?: The Pragmatic Dimensions of Genetic Explanations. Biol. Philos. 14: 349–373

Gibson TJ, Seiler M & Veitia RA (2013) The transience of transient overexpression. Nat. Methods Available at: https://www.nature.com/articles/nmeth.2534 [Accessed April 24, 2018]

Gifford F (1990) Genetic traits. Biol. Philos. 5: 327–347

Gould SJ & Lewontin RC (1979) The spandrels of San Marco and the Panglossian paradigm: a critique of the adaptationist programme. Proc R Soc Lond B 205: 581–598

Hao Y, Colak R, Teyra J, Corbi-Verge C, Ignatchenko A, Hahne H, Wilhelm M, Kuster B, Braun P, Kaida D, Kislinger T & Kim PM (2015) Semi-supervised Learning Predicts Approximately One Third of the Alternative Splicing Isoforms as Functional Proteins. Cell Rep. 12: 183–189

Hsu S-N & Hertel KJ (2009) Spliceosomes walk the line: splicing errors and their impact on cellular function. RNA Biol. 6: 526

Hu J, Boritz E, Wylie W & Douek DC (2017) Stochastic principles governing alternative splicing of RNA. PLOS Comput. Biol. 13: e1005761

Hyung D, Kim J, Cho SY & Park C (2018) ASpedia: a comprehensive encyclopedia of human alternative splicing. Nucleic Acids Res. 46: D58–D63

Kelemen O, Convertini P, Zhang Z, Wen Y, Shen M, Falaleeva M & Stamm S (2013) Function of alternative splicing. Gene 514: 1–30

Kennedy D (2004) The old file-drawer problem. Science 305: 451

Kopp F & Mendell JT (2018) Functional Classification and Experimental Dissection of Long Noncoding RNAs. Cell 172: 393–407

Kovacs E, Tompa P, Liliom K & Kalmar L (2010) Dual coding in alternative reading frames correlates with intrinsic protein disorder. Proc. Natl. Acad. Sci. U. S. A. 107: 5429–5434

Kriventseva EV, Koch I, Apweiler R, Vingron M, Bork P, Gelfand MS & Sunyaev S (2003) Increase of functional diversity by alternative splicing. Trends Genet. 19: 124–128

Kurmangaliyev YZ & Gelfand MS (2008) Computational analysis of splicing errors and mutations in human transcripts. BMC Genomics 9: 13

Light S & Elofsson A (2013) The impact of splicing on protein domain architecture. Curr. Opin. Struct. Biol. 23: 451–458

Lipscombe D, Andrade A & Allen SE (2013) Alternative splicing: functional diversity among voltage-gated calcium channels and behavioral consequences. Biochim. Biophys. Acta 1828: 1522–1529

Lipscombe D, Pan JQ & Gray AC (2002) Functional diversity in neuronal voltage-gated calcium channels by alternative splicing of Ca(v)alpha1. Mol. Neurobiol. 26: 21–44

Lonsdale J, Thomas J, Salvatore M, Phillips R, Lo E, Shad S, Hasz R, Walters G, Garcia F, Young N, Foster B, Moser M, Karasik E, Gillard B, Ramsey K, Sullivan S, Bridge J, Magazine H, Syron J, Fleming J, et al (2013) The Genotype-Tissue Expression (GTEx) project. Nat. Genet. 45: 580–585

Ma X, Kawamoto S, Uribe J & Adelstein RS (2006) Function of the neuron-specific alternatively spliced isoforms of nonmuscle myosin II-B during mouse brain development. Mol. Biol. Cell 17: 2138–2149

Melamud E & Moult J (2009) Stochastic noise in splicing machinery. Nucleic Acids Res. 37: 4873–4886

Mudge JM, Frankish A, Fernandez-Banet J, Alioto T, Derrien T, Howald C, Reymond A, Guigó R, Hubbard T & Harrow J (2011) The origins, evolution, and functional potential of alternative splicing in vertebrates. Mol. Biol. Evol. 28: 2949–2959

Pan Q, Shai O, Lee LJ, Frey BJ & Blencowe BJ (2008) Deep surveying of alternative splicing complexity in the human transcriptome by high-throughput sequencing. Nat. Genet. 40: 1413–1415

Pickrell JK, Pai AA, Gilad Y & Pritchard JK (2010) Noisy Splicing Drives mRNA Isoform Diversity in Human Cells. PLOS Genet 6: e1001236

Ramanouskaya TV & Grinev VV (2017) The determinants of alternative RNA splicing in human cells. Mol. Genet. Genomics: 1–21

Reyes A, Anders S, Weatheritt RJ, Gibson TJ, Steinmetz LM & Huber W (2013) Drift and conservation of differential exon usage across tissues in primate species. Proc. Natl. Acad. Sci. 110: 15377–15382

Rodriguez JM, Rodriguez-Rivas J, Di Domenico T, Vázquez J, Valencia A & Tress ML (2018) APPRIS 2017: principal isoforms for multiple gene sets. Nucleic Acids Res. Available at: https://academic.oup.com/nar/article/doi/10.1093/nar/gkx997/4561658/APPRIS-2017-principal-isoforms-for-multiple-gene [Accessed October 23, 2017]

Saudemont B, Popa A, Parmley JL, Rocher V, Blugeon C, Necsulea A, Meyer E & Duret L (2017) The fitness cost of mis-splicing is the main determinant of alternative splicing patterns. Genome Biol. 18: 208

Scotton P, Bleckmann D, Stebler M, Sciandra F, Brancaccio A, Meier T, Stetefeld J & Ruegg MA (2006) Activation of muscle-specific receptor tyrosine kinase and binding to dystroglycan are regulated by alternative mRNA splicing of agrin. J. Biol. Chem. 281: 36835–36845

Shehu A, Barbará D & Molloy K (2016) A Survey of Computational Methods for Protein Function Prediction. In Big Data Analytics in Genomics, Wong K-C (ed) pp 225–298. Springer International Publishing Available at: http://link.springer.com/chapter/10.1007/978-3-319-41279-5_7 [Accessed October 31, 2016]

Sievers F, Wilm A, Dineen D, Gibson TJ, Karplus K, Li W, Lopez R, McWilliam H, Remmert M, Söding J, Thompson JD & Higgins DG (2011) Fast, scalable generation of high-quality protein multiple sequence alignments using Clustal Omega. Mol. Syst. Biol. 7: 539

Smedley D, Haider S, Ballester B, Holland R, London D, Thorisson G & Kasprzyk A (2009) BioMart-biological queries made easy. BMC Genomics 10: 22

Stamm S, Ben-Ari S, Rafalska I, Tang Y, Zhang Z, Toiber D, Thanaraj TA & Soreq H (2005) Function of alternative splicing. Gene 344: 1–20

Stetefeld J & Ruegg MA (2005) Structural and functional diversity generated by alternative mRNA splicing. Trends Biochem. Sci. 30: 515–521

Tress ML, Abascal F & Valencia A (2017a) Most Alternative Isoforms Are Not Functionally Important. Trends Biochem. Sci. Available at: http://www.sciencedirect.com/science/article/pii/S0968000417300713 [Accessed May 9, 2017]

Tress ML, Abascal F & Valencia A (2017b) Alternative Splicing May Not Be the Key to Proteome Complexity. Trends Biochem. Sci. 42: 98–110

Wang ET, Sandberg R, Luo S, Khrebtukova I, Zhang L, Mayr C, Kingsmore SF, Schroth GP & Burge CB (2008a) Alternative isoform regulation in human tissue transcriptomes. Nature 456: 470–476

Wang ET, Sandberg R, Luo S, Khrebtukova I, Zhang L, Mayr C, Kingsmore SF, Schroth GP & Burge CB (2008b) Alternative isoform regulation in human tissue transcriptomes. Nature 456: 470–476

Wang M, Zhang P, Shu Y, Yuan F, Zhang Y, Zhou Y, Jiang M, Zhu Y, Hu L, Kong X & Zhang Z (2014) Alternative splicing at GYNNGY 5’ splice sites: more noise, less regulation. Nucleic Acids Res. 42: 13969–13980

Worton LE, Shi Y-C, Smith EJ, Barry SC, Gonda TJ, Whitehead JP & Gardiner EM (2017) Ectodermal-Neural Cortex 1 Isoforms Have Contrasting Effects on MC3T3-E1 Osteoblast Mineralization and Gene Expression. J. Cell. Biochem. 118: 2141–2150

Zhang Z, Xin D, Wang P, Zhou L, Hu L, Kong X & Hurst LD (2009) Noisy splicing, more than expression regulation, explains why some exons are subject to nonsense-mediated mRNA decay. BMC Biol. 7: 23

Zhao S & Zhang B (2015) A comprehensive evaluation of ensembl, RefSeq, and UCSC annotations in the context of RNA-seq read mapping and gene quantification. BMC Genomics 16: 97

